# FKBP11 targeted plasma cells promotes abdominal aortic aneurysm progression through an m6A-dependent mechanism

**DOI:** 10.1101/2024.05.05.592616

**Authors:** Yuchen He, Jia Xing, Shiyue Wang, Han Jiang, Yu Lun, Yanshuo Han, Philipp Erhart, Böckler Dittmar, Jian Zhang

## Abstract

**Objective:** Despite surgical advance, effective targeted drugs for non-surgical treatment of abdominal aortic aneurysm (AAA) are lacking because of the unclear pathogenesis of AAA. N6-methyladenosine (m6A) methylation, acknowledged for its pivotal influence on RNA metabolism, including aspects such as stability, transport, translation, and splicing, is largely implied for its role in AAA mechanism. This study aims to elucidate the involvement of m6A methylation in the progression of AAA through an integrative multi-omics and machine learning approach.

**Methods and Results:** We utilized methylated RNA immunoprecipitation sequencing (MeRIP-seq) to map the m6A methylation landscape in AAA tissues and combined this with RNA sequencing (RNA-seq) from the GEO database, to explore the interplay between m6A methylation and gene expression. A machine learning-based AAA m6A-related mRNA signature (AMRMS) was developed to predict the risk of AAA dilation. The AMRMS showed robust predictive power in distinguishing between patients with large and small AAAs. Notably, FKBP11 was identified as a key gene significantly influencing the predictive model, and up-regulated in large AAAs compared to its in small AAAs. Further single-cell RNA sequencing (scRNA-seq) and histological analysis highlighted the critical role of FKBP11 in mediating the endoplasmic reticulum stress of plasma cells within the AAA walls and its correlation with m6A methylation.

**Conclusions:** The m6A modification regulatory network plays a vital role in the progression of AAA, and the AMRMS offers promising potential in assessing the risk of AAA dilation. Our findings suggest that elevated FKBP11, by activating endoplasmic reticulum stress in plasma cells, may significantly contribute to AAA expansion.

## Introduction

Abdominal aortic aneurysm (AAA), a common degenerative artery dilatory disease in the elderly, affects about 1%-2% of this group [1, 2]. Many individuals with AAA show no symptom but sudden rupture, leading to a life-threatening complication, highlighting the unpredictability of this result [1]. Despite advancements in surgery such as open surgery and endovascular repair which offer positive outcomes for patients whose aneurysms meet the criteria for surgery, a lack of understanding in the mechanism from occurrence to development of AAA leaves those non-operation indicated patients without targeted medical treatments to halt disease progression. Understanding these mechanisms is crucial for the creation of novel diagnostic approaches to evaluate and predict AAA progression and novel therapeutic approaches to improve patient prognosis.

The role of M6A methylation, marked by the methylation of adenosine nucleotides within RNA molecules, is increasingly acknowledged for its pivotal influence on RNA metabolism, including aspects such as stability, transport, translation, and splicing [3]. Dynamic regulation of RNA by m6A methylation has been proven to facilitate crucial pathological transformations [4], including apoptosis, remodeling of the extracellular matrix, and infiltration by inflammatory cells, the processes deeply entwined with the pathogenesis and progression of AAA [5–7]. Notably, our research team has identified the presence of m6A methylation modifications throughout the development of AAA, and underscored its significance [8]. Yet, the complex regulatory network of m6A methylation and the heterogeneous cellular composition of the AAA vessel wall pose challenges in fully understanding m6A’s role in the progression of AAA.

In this study, we conducted a comprehensive analysis of the m6A methylome in AAA tissue samples using Methylated RNA Immunoprecipitation Sequencing (MeRIP-seq), thereby providing a transcriptome-wide landscape of m6A modifications within AAA tissues. This was integrated with RNA sequencing to explore the relationship between m6A methylation levels and gene expressions; while machine learning analyses were employed to underscore the significance of m6A methylation in predicting the progression from small to large aneurysms. Furthermore, single cell RNA sequencing (scRNA-seq) was utilized to link m6A methylation with the complex cellular components of the AAA vessel wall. Through a multidimensional, multi-omics approach, we elucidate the complex role of the m6A regulatory network in the AAA development.

## Methods

The flow chart of the current research was shown in **Figure 1**.

**Figure 1.**
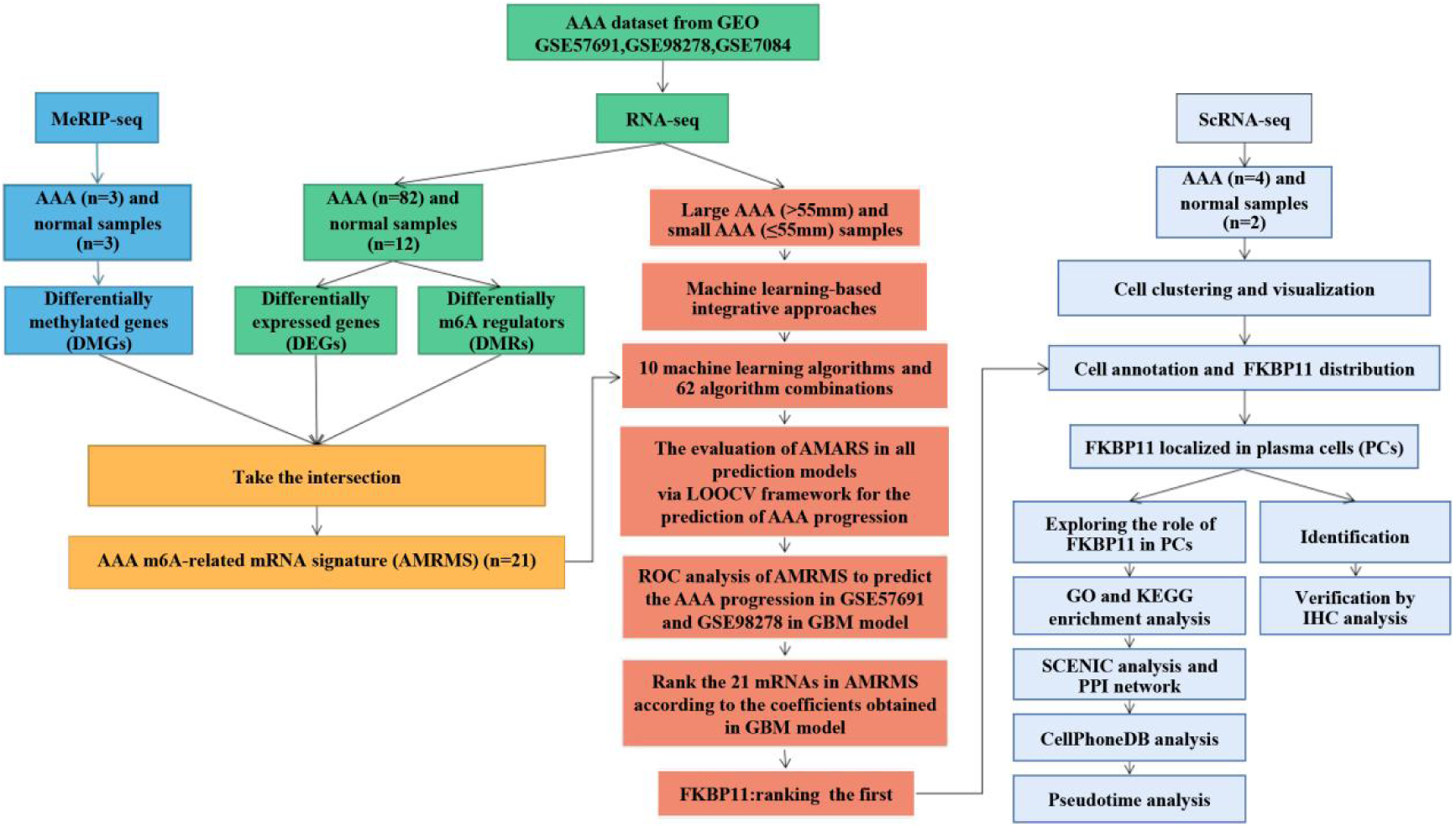
Flow chart of the research process.

### Patients and samples

Seven patients with atherosclerotic AAA were consecutively enrolled in this study. Tissue specimens from the AAA patients and their corresponding clinical data were collected after aneurysmal open surgery repair at the First Hospital of China Medical University. Patients with Ehlers-Danlos syndrome, Marfan syndrome, other known vascular or connective tissue disorders, cancer, infection, and any other immune-related disease were excluded. Among the collected AAA tissue samples, three were used for MeRIP-seq, and the remaining four samples were used for further scRNA-seq.

Simultaneously, seven control aortic tissue samples were obtained from organ donors, serving as the normal aorta control group (three for MeRIP-seq and two for scRNA-seq). These control samples had a relatively healthy peripheral vascular system, determined by Computed Tomography Angiography (CTA), and no evidence or medical history of aneurysm or other vascular disorders. For MeRIP-seq, AAA tissue samples and matched normal tissues were promptly collected and separated into centrifuge tubes. For scRNA-seq, AAA tissue samples and control samples were stored in tissue preservation solution purchased from SeekGene BioSciences Co., Ltd., Beijing, China, and transferred under 4℃ to the SeekGene BioSciences Co. laboratory. This study was approved by the Ethics Committee of China Medical University (AF-SOP-07-1.1-01).

### RNA extraction, MeRIP experiment, library preparation, and sequencing

MeRIP experiments, high-throughput sequencing and data analysis were conducted by Seqhealth Technology Co., Ltd. (Wuhan, China). Total RNAs were extracted from three AAA samples and three healthy control (HC) samples using TRIzol Reagent (Invitrogen, United States) following the manufacturer’s protocol. Subsequently, DNA digestion was carried out after RNA extraction using DNaseI. RNA quality was assessed by examining A260/A280 with a Nanodrop spectrophotometer (Thermo Fisher Scientific Inc., United States), and RNA integrity was confirmed by 1.5% agarose gel electrophoresis. Qualified RNAs were quantified using Qubit3.0 with the QubitTM RNA Broad Range Assay kit (Life Technologies, United States).

For polyadenylated RNA enrichment, 50µg of total RNAs was used with VAHTS mRNA Capture Beads (Vazyme, China). RNA fragments were mainly distributed in the 100-200 nt range after the addition of 20mM ZnCl2 and incubation at 95℃ for 5-10 minutes. Then, 10% of RNA fragments were saved as “Input,” and the remaining fragments were used for m6A immunoprecipitation (IP) using a specific anti-m6A antibody (Synaptic Systems, 202203, Germany). RNA samples from both input and IP were prepared using TRIzol reagent (Invitrogen). The stranded RNA sequencing library was constructed using the KC-DigitalTM Stranded mRNA Library Prep Kit for Illumina (Seqhealth, China) following the manufacturer’s instructions. The kit employed unique molecular identifiers (UMI) of 8 random bases to label pre-amplified cDNA molecules, thereby eliminating duplication bias in PCR and sequencing steps. The library products, corresponding to 200-500 bps, were enriched, quantified, and finally sequenced on a Novaseq 6000 sequencer (Illumina) with PE150 model.

### Data analysis for MeRIP-seq

Raw sequencing data were initially filtered by Trimmomatic (version 0.36) to discard low-quality reads and trim reads contaminated with adaptor sequences. Clean reads underwent further processing with in-house scripts to eliminate duplication bias introduced during library preparation and sequencing. Clean reads were clustered based on UMI sequences, grouping reads with the same UMI sequence together. Within each cluster, pairwise alignment was performed to identify reads with sequence identities exceeding 95%, which were then extracted into new sub-clusters. Multiple sequence alignment was conducted to obtain a consensus sequence for each sub-cluster, eliminating errors and biases introduced by PCR amplification or sequencing.

The de-duplicated consensus sequences were used for m6A site analysis, mapping to the reference genome of Homo sapiens from NCBI using STAR software (Version 2.5.3a) with default parameters. The exomePeak (Version 3.8) software was employed for peak calling, and m6A peaks were annotated using bedtools (Version 2.25.0). Differentially methylated peaks were identified using a python script with the Fisher test. Sequence motifs enriched in m6A peak regions were determined using Homer (version 4.10). Differentially methylated genes (DMGs) and specific methylated genes (SMGs) were identified using the corresponding MeRIP-seq input library data by the R package “Ballgown”. Gene Ontology (GO) analysis and Kyoto Encyclopedia of Genes and Genomes (KEGG) enrichment analysis were performed based on annotated genes using KOBAS software (version: 2.1.1) with a corrected p value cutoff of 0.05 for statistically significant enrichment.

### Public databases and analysis

Three mRNA sequencing datasets of humans were downloaded from the GEO database (https://www.ncbi.nlm.nih.gov/geo/), including GSE57691, GSE7084, and GSE98278. A total of 82 AAA tissue samples and 12 normal aortic tissue samples were selected and further normalized and integrated for analysis. Principal Component Analysis (PCA) was conducted using the “mixOmics” R package to detect the independence of the AAA group and the control group. The stromal score and immune score differences between the two groups were evaluated using the “Estimate” R package. Based on Wilcoxon test method, differential Expressed Genes (DEGs) between the two groups were identified with a statistical threshold of |log2Fold Change (FC)| > 1 and P < 0.05 using the “limma” R package. DEGs and DMGs were then overlapped, and the intersecting genes were considered as hub genes. A heatmap diagram was constructed to display the expression of hub genes in AAA and normal aortic tissues.

Currently, twenty-one genes, including ALKBH5, FTO, HNRNPA2B1, HNRNPC, IGF2BP1, IGF2BP2, IGF2BP3, FMR1 METTL14, METTL3, RBM15, RBM15B, CBLL1, WTAP, YTHDC1, YTHDC2, YTHDF1, YTHDF2, YTHDF3, ZC3H13, and ELAVL1, are recognized as common m6A RNA methylation regulators [9]. A boxplot was generated using the “ggpubr” R package to visualize the expression of these m6A regulators in the two groups.

### Establishment of AAA m6A-related mRNA signature (AMRMS) and assessment of the robustness of signature in a machine learning-based approaches

To develop a consensus AMRMS with high accuracy and stability performance, we integrated 12 machine learning algorithms and 101 algorithm combinations to fit prediction models. The GSE57691 dataset was regarded as training set, while GSE98278 dataset was for validation set. Twelve machine learning algorithms were incorporated, including Lasso, plsRglm (partial least squares regression generalized linear model), Elastic Network (Enet), Ridge, Generalized Boosted Regression Modeling (GBM), Support Vector Machine (SVM), Random Forest (RF), Linear Discriminant Analysis (LDA), Generalized Linear Model Boosting (glmBoost), eXtreme Gradient Boosting (XGBoost), Naive Bayes, and Step Generalized Linear Model (Stepglm). Based on the leave-one-out cross-validation (LOOCV) framework in the training set, all models were detected across the validation dataset, and the model with the highest average Harrell’s concordance index (C-index) was considered optimal. The algorithm’s prediction ability was measured using the area under curve (AUC), and finally the combination of algorithms with both robust performance and clinically prediction significance was selected. According, we established a final signature that can predict the robustness of AAA patients, called AAA m6A-related mRNA signature (AMRMS).

### The construction and validation of AMRMSscore model

The AMRMS of each sample was quantitatively analyzed by single sample gene set enrichment analysis (ssGSEA), and we further obtained the AMRMSscore for each patient. According to the median ssGSEA score, patients were divided into two groups: low risk group and high-risk group, and the expressions of genes enrolled in AMRMS between these two group were visualized by violin plot.

### Cell preparation for scRNA-seq

After harvesting, tissues were washed in ice-col d RPMI1640 and dissociated using the multi-tissue dissociation kit 2 (Miltenyi, Germany) according to the manufacturer’s instructions. DNase treatment was optional, depending on the viscosity of the homogenate. Cell count and viability were estimated using a fluorescence Cell Analyzer (Countstar Rigel S2) with AO/PI reagent after removing erythrocytes (Miltenyi) and any debris and dead cells were removed if necessary (Miltenyi). Finally, fresh cells were washed twice in RPMI1640 and resuspended at 1×10^6^ cells/ml in 1×PBS and 0.04% bovine serum albumin.

### ScRNA-seq library construction and sequencing

In this study, we utilized SeekGene’s SeekOne® MM Single Cell 3’ library preparation kit to construct scRNA-seq libraries with improved efficiency and reduced duplication rate. We began by loading the appropriate number of cells into the flow channel of the SeekOne® MM chip, containing 170,000 microwells, using gravity to aid cell settling. After allowing sufficient time for settling, any remaining unsettled cells were carefully removed.

To label individual cells, we employed Cell Barcoded Magnetic Beads (CBBs), which were pipetted into the flow channel and precisely localized within the microwells using a magnetic field. Subsequently, we lysed the cells within the SeekOne® MM chip, releasing RNA that was captured by the CBBs in the same microwell. Reverse transcription was performed at 37℃ for 30 minutes to synthesize cDNA with cell barcodes. Exonuclease I treatment was used to eliminate any unused primers on the CBBs. The barcoded cDNA on the CBBs then underwent hybridization with a random primer containing a Reads 2 SeqPrimer sequence at its 5’ end, facilitating the extension and creation of the second DNA strand with cell barcodes at the 3’ end. After denaturation, the resulting second strand DNA was separated from the CBBs and purified. The purified cDNA product was amplified through PCR, and unwanted fragments were removed through a clean-up process. Full-length sequencing adapters and sample indexes were added to the cDNA through an indexed PCR. Indexed sequencing libraries were further purified using SPRI beads to remove any remaining impurities. Quantification of the libraries was performed using quantitative PCR (KAPA Biosystems KK4824). Finally, the libraries were sequenced using either the Illumina NovaSeq 6000 platform with PE150 read length or the DNBSEQ-T7 platform with PE100 read length.

### ScRNA-seq data quality control, batch effect correction, and clustering scRNA-seq data processing

By using the “Seurat” package, the low-quality cells of scRNA-seq data were first excluded with more than 25% mitochondrial gene, and genes expressed in at least three cells within an expression range of 100 to 4000. We then identified the highly variable genes were calculated using the ‘vst’ method, and the scaled data were then removed batch effect from six samples by “Harmony” package. Cell clusters were constructed by using the “FindClusters” and “FindNeighbors” functions, and then clusters were projected to t-SNE and UMAP.

### Cell annotation analysis

Cell types of different clusters were identified manually using cluster-specific marker genes from databases and literatures. To employed to explore the cellular landscape and gene expression, various visualization techniques, including UMAP, t-SNE, dotplot, and featureplot were utilized.

### DEG and enrichment analysis for scRNA-seq

The FindMarkers function was used to identify DEGs across the conditions using the default Wilcoxon test. Genes were ranked by absolute log2FC, and those with P values > 0.05 (adjusted for multiple comparisons) and log2FC < 0.25 were removed.

GO enrichment analysis of DEGs and highly variable genes was implemented using the “clusterProfiler” R package. GO terms with corrected P-values less than 0.05 were considered significantly enriched by DEGs. The “clusterProfiler” R package was also used to test the statistical enrichment of DEGs in KEGG pathways.

Based on the Hallmark genesets obtained from Molecular Signatures Database, we used the “area under the curve” (AUC) method to calculate the AUCell scores of single cells based on the input gene signature. A higher AUCell score in a single cell represents a more enriched activity of the gene set of interest. An adjusted P <0.05 was considered statistically enriched.

### Assessment of transcription factor (TF) activity

To perform transcription factor analysis of each cell subgroup, we utilized VIPER in combination with DoRothEA to calculate their TF activity scores and then used heatmap for visualization.

### Analysis of signaling pathway activity

We applied PROGENy to estimate the signaling pathway activity of each cell type, and all parameters were set to default values. The activity of top 30 signailing pathways was shown by heatmap.

### The analysis of the differentiation potential of different single cell subpopulations

We performed CytoTRACE analysis to predict differentiation states of different subclusters, and complement the trajectory analysis from Monocle. The top correlated genes were also analyzed and visualized based on CytoTRACE algorithm.

### Pseudotime analysis

Single-cell pseudotime trajectory analysis was performed with the “Monocle2” package to determine the potential lineage differentiation of B cell subclusters. Based on DEGs among Seurat clusters, cell progression genes were defined, and DifferentialGeneTest function was further used to explore gene dynamics during cell differentiation. We finally used Monocle to map out the evolutionary trajectory using a reverse graph embedding algorithm, superimposed the trajectory onto a UMAP plot, and assigned a pseudo-time value to each cell subclusters. Additionally, the changing trend of focus genes with cell differentiation was revealed by evaluating the gene expression of cells in each segment.

### Immunohistochemistry (IHC) analysis

IHC analysis was performed to investigate the expression patterns of specific biomarkers within the aortic tissue samples. Representative sections of the samples from CMU Aneurysm Biobank (CMUaB), measuring 2–3 μm in thickness, were utilized for this analysis. Consecutive slides from each specimen were meticulously prepared and subjected to incubation with appropriate antibodies to precisely determine the expression of the biomarkers in individual cells within the aneurysmal lesions.

In each case, one slide was stained using an antibody to detect the specific cell type present, while a consecutive slide was stained using the antibody targeting the individual biomarker of interest. Paraffin sections were routinely stained for IHC analysis, following well-established protocols as previously described [8].

The primary antibodies employed in this study included an anti-J Chain antibody (dilution 1:5000; Husbio, Wuhan, China), utilized for the detection of plasma cells, and an anti-FKBP11 antibody (dilution 1:200; Absin, Shanghai, China), used to specifically target FKBP11 expression.

### Transmission electron microscopy (TEM)

The fresh AAA samples obtained through elective open surgical repair was applied for the analysis of TEM. The relative protocols have been previously described [10]. The positive cells were examined by TEM (JEM-1200EX, JEOL, Tokyo, Japan).

### Statistical analysis

All statistical analyses were performed using the R language. Continuous variables between the 2 groups were compared using the Wilcoxon-rank sum test, while the correlation was evaluate by the Spearman correlation analysis. A P value <0.05 was considered statistically significant in all analyses.

## Results

### Comparative m6A methylation patterns in AAA and normal aortic tissue

Our study presents a detailed MeRIP-seq analysis that explores the m6A methylation landscape across human AAA and healthy aortic tissues. The participant baseline data is outlined in **Table 1**. We identified a differential m6A methylome with 6988 and 7256 m6A peaks in AAA and normal control (NC) tissues, respectively (**Figure 2A**). Of these, 5947 were shared, while 1309 were unique to AAA and 1041 unique to NC, as depicted in the Venn diagram (**Figure 2B**).

**Figure 2.**
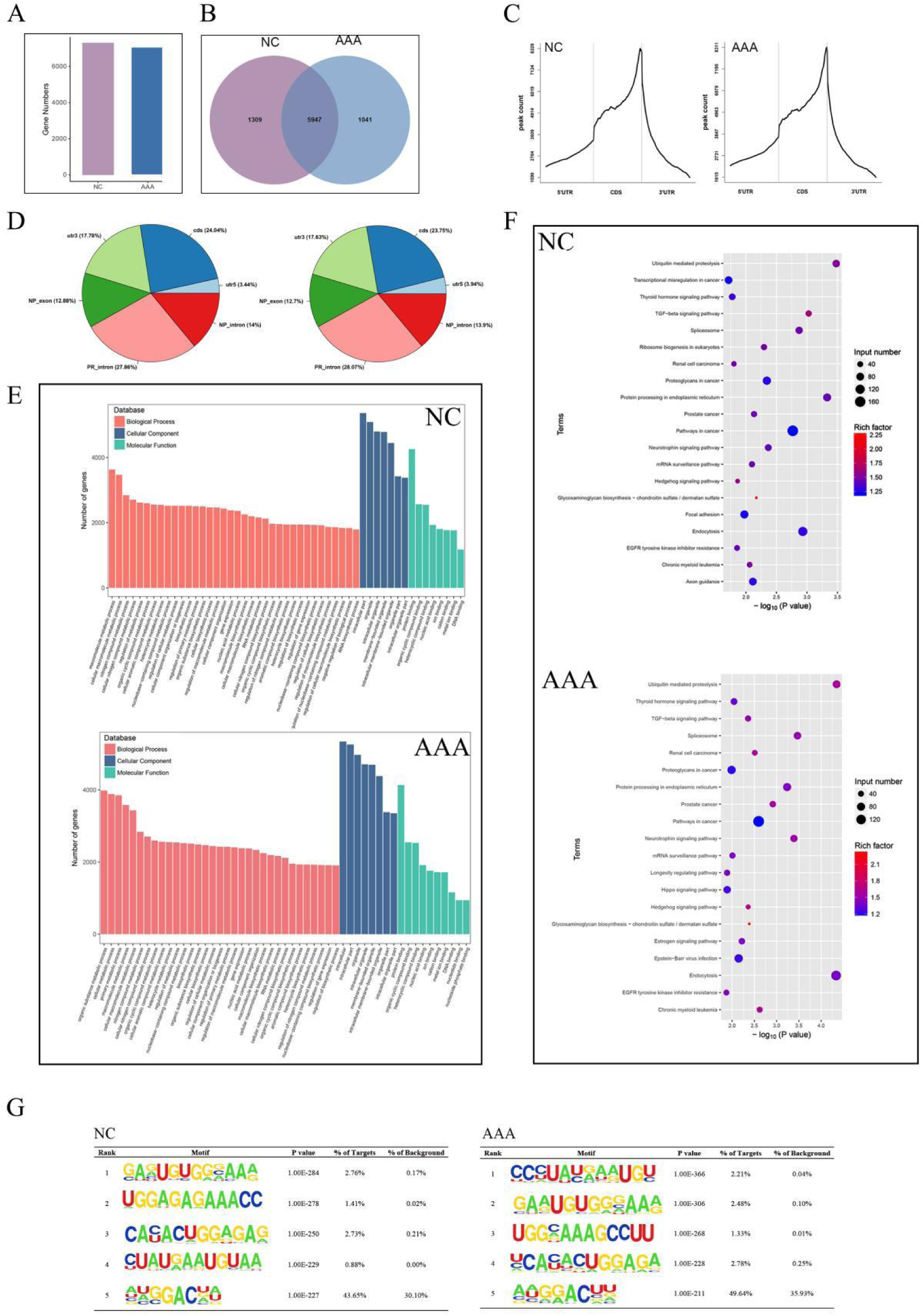
Transcriptome-wide MeRIP-seq and analysis of m6A peaks. (**A**) Overlap of m6A peaks in the AAA group and the normal control (NC) group. (**B**) Numbers of AAA-unique, NC-unique, and common m6A peaks are shown as Venn diagram as well as m6A peaks representing genes in the two groups. (**C**) and (**D**) Proportion of m6A peaks distributed in the indicated regions in the NC group and the AAA. (**E**) Major gene ontology terms significantly enriched in the two groups. (**F**) Bubble plots of major KEGG terms significantly enriched in the two groups. (**G**) Top 5 m6A modification motifs enriched from all identified m6A peaks in the two groups.

**Table 1.**
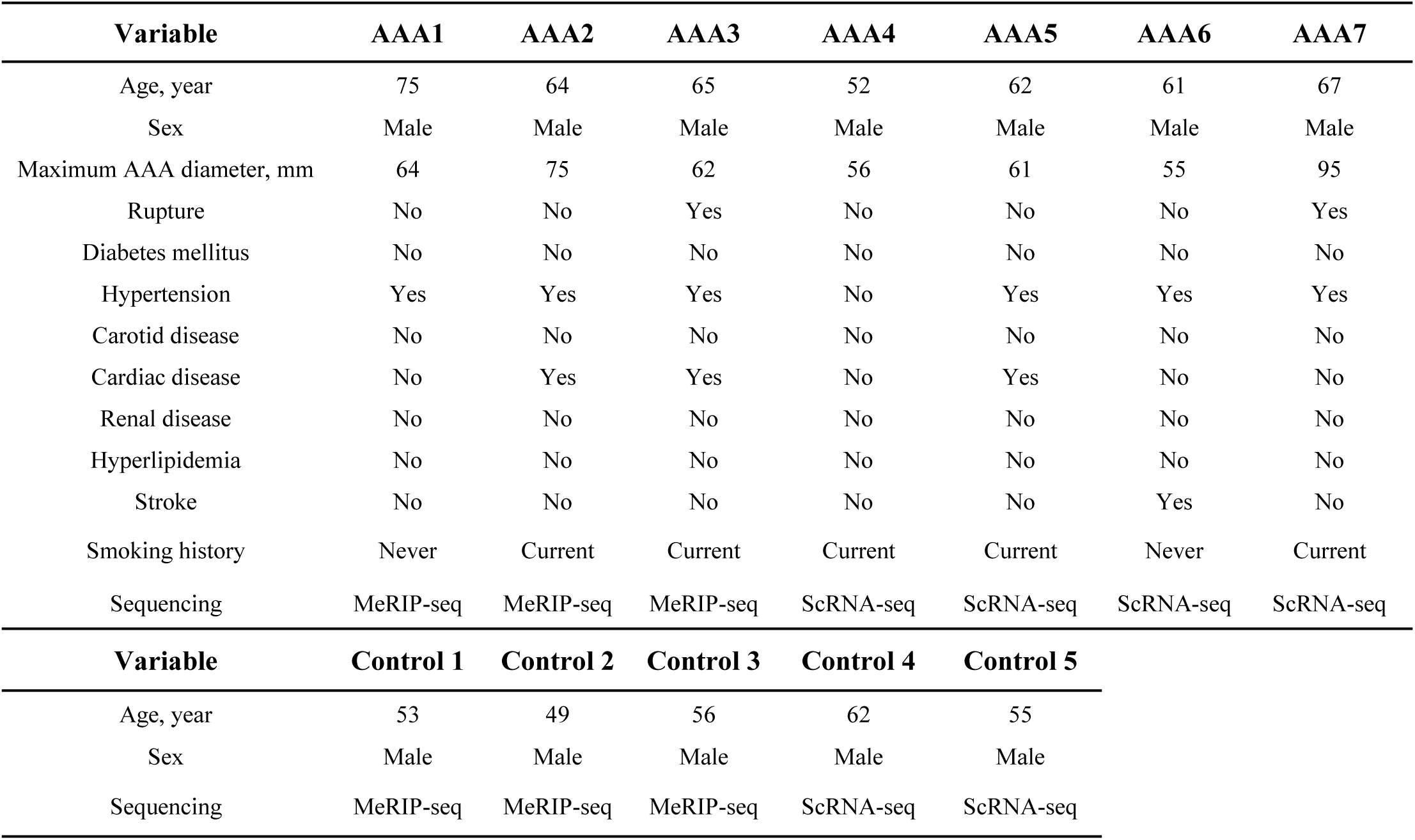
The baseline data of AAA patients (n=10) included in this study.

| Table 1 The baseline data of AAA patients (n=10) included in this study |  |  |  |  |  |  |  |
| --- | --- | --- | --- | --- | --- | --- | --- |
| Variable | AAA1 | AAA2 | AAA3 | AAA4 | AAA5 | AAA6 | AAA7 |
| Age, year | 75 | 64 | 65 | 52 | 62 | 61 | 67 |
| Sex | Male | Male | Male | Male | Male | Male | Male |
| Maximum AAA diameter, mm | 64 | 75 | 62 | 56 | 61 | 55 | 95 |
| Rupture | No | No | Yes | No | No | No | Yes |
| Diabetes mellitus | No | No | No | No | No | No | No |
| Hypertension | Yes | Yes | Yes | No | Yes | Yes | Yes |
| Carotid disease | No | No | No | No | No | No | No |
| Cardiac disease | No | Yes | Yes | No | Yes | No | No |
| Renal disease | No | No | No | No | No | No | No |
| Hyperlipidemia | No | No | No | No | No | No | No |
| Stroke | No | No | No | No | No | Yes | No |
| Smoking history | Never | Current | Current | Current | Current | Never | Current |
| Sequencing | MeRIP-seq | MeRIP-seq | MeRIP-seq | ScRNA-seq | ScRNA-seq | ScRNA-seq | ScRNA-seq |
| Variable | Control 1 | Control 2 | Control 3 | Control 4 | Control 5 |  |  |
| Age, year | 53 | 49 | 56 | 62 | 55 |  |  |
| Sex | Male | Male | Male | Male | Male |  |  |
| Sequencing | MeRIP-seq | MeRIP-seq | MeRIP-seq | ScRNA-seq | ScRNA-seq |  |  |

The distribution analysis of m6A sites highlighted a concentration in the coding sequences (CDS), particularly adjacent to the 3’UTR near stop codons **(Figures 2C** and **2D**). Interestingly, we observed a similar distribution pattern in CDS, 3’UTR, and 5 ‘UTR regions between AAA and healthy aorta, underscoring conserved m6A deposition in these transcript zones.

### Functional implications of m6A methylation in aortic tissues

GO and KEGG pathway analyses were conducted on genes with m6A modifications, revealing their involvement in various biological processes and pathways (**Figures 2E** and **2F**). These findings were integral in deciphering the biological significance of m6A modifications in both healthy and diseased aortic environments.

We further characterized the top five m6A motifs enriched in the m6A peaks of both groups, illuminating potential regulatory motifs integral to the epigenetic mechanisms of AAA (**Figure 2G**).

### Altered m6A peak analysis reveals differentially methylated mRNAs

In the comparative analysis, we observed 805 m6A peaks up-regulated and 559 peaks down-regulated in AAA versus NC (| log2FC | > 1 and p < 0.05). The top 20 differentially regulated m6A peaks for both up- and down-regulated transcripts are enumerated in **Tables 2** and **Table 3**, respectively.

**Table 2.** Top 20 up-regulated genes in the AAA group compared with the NC group.

| Gene | Chromosome | Peak Start | Peak End | logFC | P Value | Annotation |
| --- | --- | --- | --- | --- | --- | --- |
| RP11-458D21.1 | chr1 | 146058976 | 146059187 | 8.549092311 | 3.96E-258 | Exon |
| AC006460.2 | chr2 | 190699764 | 190701481 | 8.331286675 | 9.26E-100 | Exon |
| ANKRD35 | chr1 | 145872259 | 145872349 | 6.572519874 | 3.46E-100 | CDS |
| RAB6B | chr3 | 133826000 | 133826120 | 6.103135181 | 2.35E-25 | 3'UTR |
| RP11-190A12.9 | chr1 | 159910150 | 159910267 | 6.091453376 | 1.31E-86 | Exon |
| ANKRD35 | chr | 145872617 | 145873155 | 5.252018306 | 2.52E-49 | CDS |
| PEX11B | chr1 | 145912398 | 145912549 | 4.918719817 | 2.67E-50 | CDS |
| CCNT2 | chr2 | 134918621 | 134918710 | 4.801015567 | 1.35E-17 | 5'UTR |
| SLC16A9 | chr10 | 59654199 | 59654409 | 4.529542873 | 1.71E-68 | CDS |
| RP11-956E11.1 | chr12 | 69459578 | 69460640 | 4.509641267 | 7.00E-33 | Exon |
| B9D1 | chr17 | 19377782 | 19377902 | 4.48020483 | 1.39E-17 | Exon |
| SH3BP5L | chr1 | 248825914 | 248826005 | 4.425779076 | 4.29E-07 | 5'UTR |
| IRS2 | chr13 | 109752906 | 109753057 | 4.382558547 | 2.26E-07 | 3'UTR |
| NRG1 | chr8 | 32764036 | 32764247 | 4.376053202 | 1.56E-15 | CDS |
| CLN5 | chr13 | 76991645 | 76991855 | 4.288490223 | 3.39E-08 | 5'UTR |
| PEG10 | chr7 | 94663468 | 94663528 | 4.231928235 | 4.23E-12 | CDS |
| SP4 | chr7 | 21429813 | 21429934 | 4.116758465 | 1.78E-12 | CDS |
| ABCC10 | chr6 | 43432629 | 43432690 | 4.103135181 | 1.77E-35 | CDS |
| OR1K1 | chr9 | 122800508 | 122800568 | 4.090045181 | 1.43E-32 | CDS |
| SYNGAP1 | chr6 | 33456473 | 33456623 | 4.06851564 | 3.63E-05 | 3'UTR |

**Table 3.** Top 20 down-regulated genes in the AAA group compared with the NC group.

| Gene | Chromosome | Peak Start | Peak End | logFC | P Value | Annotation |
| --- | --- | --- | --- | --- | --- | --- |
| AGTR1 | chr3 | 148742109 | 148742348 | -6.336600727 | 2.23E-157 | 3'UTR |
| C1GALT1C1L | chr2 | 43675589 | 43675824 | -5.829866907 | 1.74E-27 | CDS |
| LINC01099 | chr4 | 177907817 | 177907936 | -5.82891394 | 7.46E-35 | Exon |
| APOB | chr2 | 21012100 | 21012221 | -5.67323491 | 4.17E-61 | CDS |
| PLCXD3 | chr5 | 41382142 | 41382382 | -5.420916738 | 1.69E-29 | CDS |
| KCNK17 | chr6 | 39299326 | 39299386 | -5.141089108 | 1.07E-39 | 3'UTR |
| SACS-AS1 | chr13 | 23428362 | 23428422 | -4.892928011 | 2.22E-32 | Exon |
| MAGEC3 | chrX | 141897265 | 141897326 | -4.672031829 | 1.13E-07 | CDS |
| COLEC10 | chr8 | 119105934 | 119106141 | -4.590041585 | 2.21E-13 | CDS |
| AC011625.1 | chr7 | 132367800 | 132367859 | -4.583511243 | 2.66E-15 | Exon |
| KCTD16 | chr5 | 144207227 | 144207378 | -4.471213059 | 7.98E-23 | CDS |
| KCNJ3 | chr2 | 154854794 | 154855273 | -4.460750379 | 4.28E-06 | CDS |
| TENM1 | chrX | 124380632 | 124380993 | -4.394137407 | 7.90E-12 | CDS |
| CDH8 | chr16 | 62021340 | 62021580 | -4.233979854 | 2.09E-07 | CDS |
| RP1-95L4.4 | chr6 | 143343173 | 143343383 | -4.213221928 | 4.34E-24 | Exon |
| LLNLF-187D8.1 | chr19 | 46428291 | 46428412 | -4.18325756 | 3.30E-03 | CDS |
| CD200 | chr3 | 112332233 | 112332410 | -4.027465659 | 1.51E-06 | 5'UTR |
| ADRA1D | chr20 | 4221409 | 4221560 | -4.014672632 | 4.27E-05 | 3'UTR |
| RP11-768F21.1 | chr12 | 119594280 | 119594340 | -3.992971755 | 3.74E-12 | Exon |
| FZD3 | chr8 | 28502835 | 28503076 | -3.988564653 | 3.57E-09 | CDS |

GO and KEGG pathway analyses for genes with altered m6A peaks unraveled their association with pivotal biological processes and signaling pathways, including the Notch signaling and autophagy (**Figures 3A** and **3B**). Notably, similar enrichments in both up- and down-regulated m6A peaks were observed for processes like positive regulation of mitophagy and molecular functions related to smooth muscle contraction.

**Figure 3.**
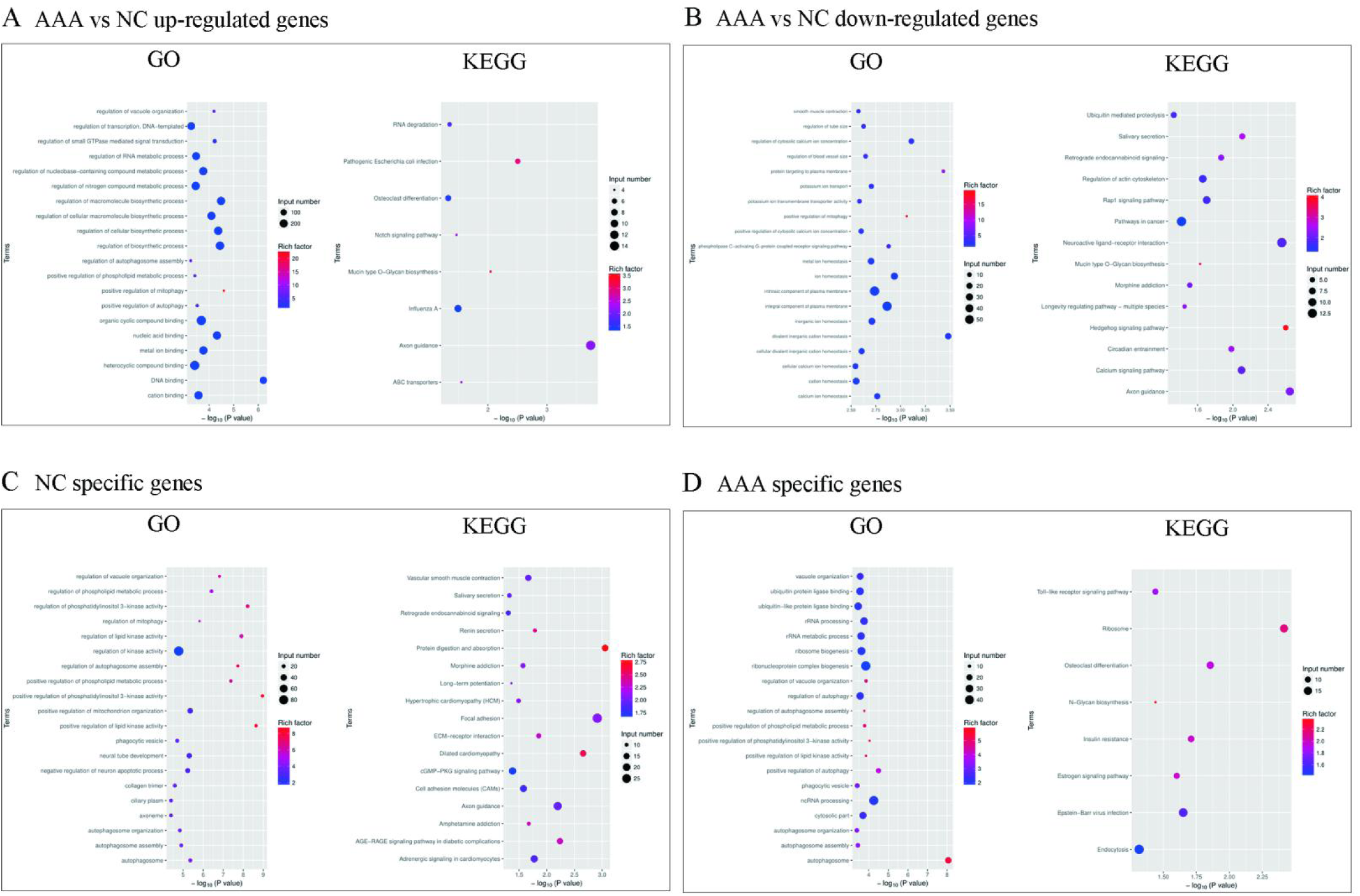
Systematic functional analysis of genes with differentially m6A modification. The Bubble plots of GO analysis and KEGG analysis for up-regulated methylated genes and down-regulated methylated genes in AAA shown in (**A**) and (**B**). Major GO enrichments and KEGG enrichments of NC specific methylated genes (**C**) and AAA specific methylated genes (**D**).

### Specific m6A peak analysis in AAA and healthy aortic tissue

When examining m6A peaks unique to AAA and NC, we identified distinct biological processes and pathways using GO and KEGG analyses (**Figures 3C** and **3D**). The AAA-specific m6A-modified genes were predominantly involved in autophagy-related processes, while NC-specific genes were associated with pathways pertinent to vascular smooth muscle activity and focal adhesion.

### Integrative analysis of MeRIP-seq and RNA-seq Data identifies key m6A regulators and hub DMGs in AAA

We performed a comprehensive analysis by integrating MeRIP-seq and RNA-seq data to identify hub DMGs in AAA. PCA clearly distinguished mRNA profiles obtained from AAA samples (n=82) and normal aortic tissue samples as controls (n=12) in the GSE57691, GSE7084, and GSE98278 datasets. To assess the differences between the two groups, we evaluated the stromal score and immune score. The results indicated that the AAA group exhibited significantly higher stromal and immune scores compared to the control group (**Figure 4A** and **4B**).

**Figure 4.**
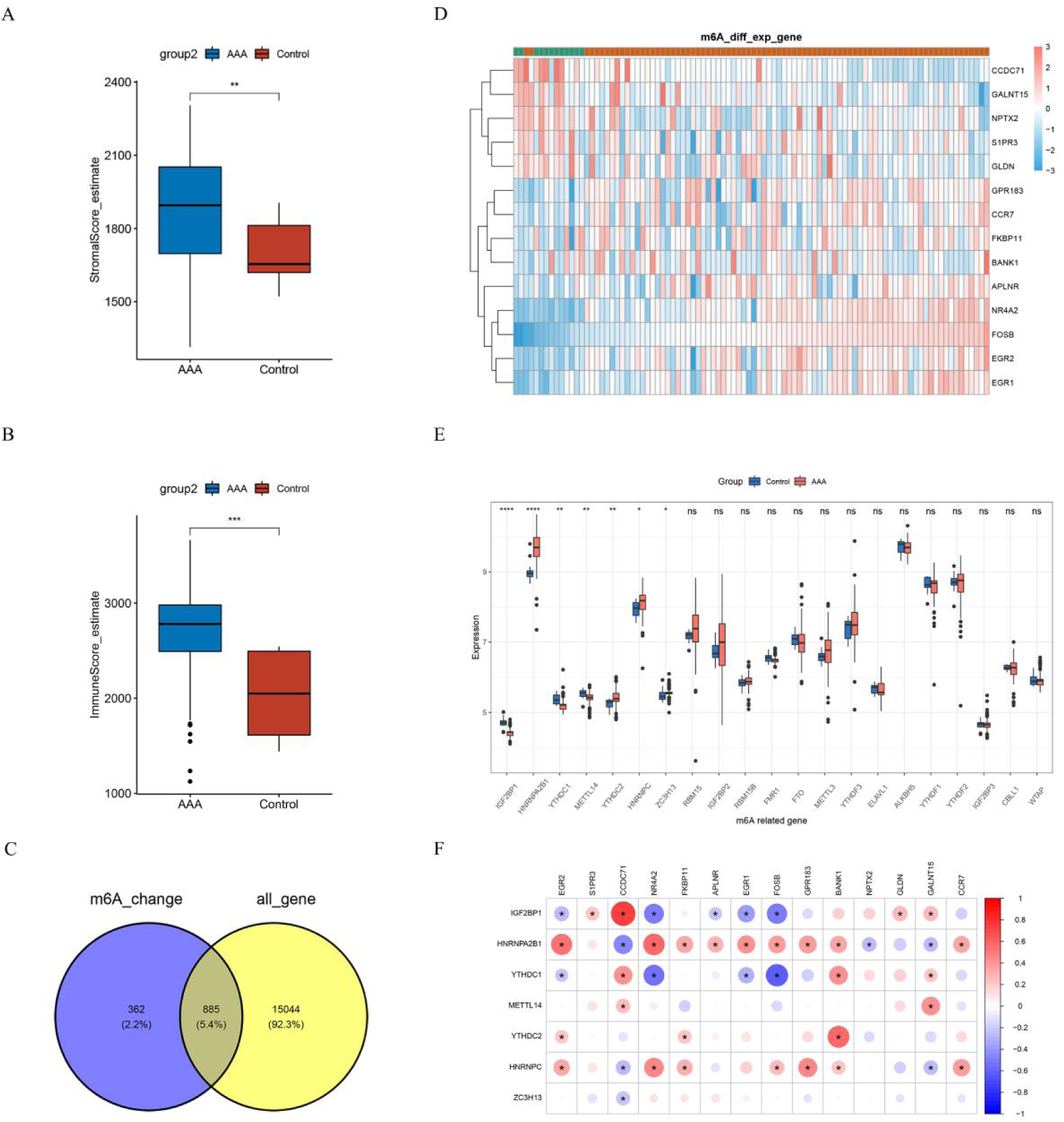
The conjoint analysis of MeRIP-seq and RNA-seq from GEO database to explore m6A expression pattern in AAA tissue samples. Comparison of stromal score (**A**) and immune score (**B**) in the AAA group and the control group shown in the histograms. (**C**) Venn diagram of DMGs from MeRIP-seq and all genes from RNA-seq. (**D**) The expressions 14of hub DMGs in 82 AAA samples and 12 control samples were shown in the heatmap. (**E**) The differential analysis of 21 m6A modulators between the AAA group and the control group was shown in the boxplots. (**F**) The correlation among hub DMGs and DMRs were shown through correlation heatmap plot.

Next, we obtained a total 885 differentially expressed DMG through overlapping all genes obtained from RNA-seq data with the DMGs identified from MeRIP-seq data (| log2FC | > 0.585 and p < 0.05). (**Figure 4C**). Then we narrowed the range of DEGs by (| log2FC | > 1 and p < 0.05). We found 14 genes that overlapped between the two datasets, which we considered as the hub DMGs (**Figure 4D** and **Table 4**). Among these hub genes, 9 genes, namely NR4A2, FKBP11, APLNR, EGR2, EGR1, FOSB, GPR183, BANK1, and CCR7, were up-regulated in the AAA group compared to the controls. On the other hand, 5 genes, including S1PR3, CCDC71, NPTX2, GLDN, and GALNT15, were down-regulated in the AAA group.

**Table 4.** Hub differentially methylated genes in AAA from the integration of GSE57691, GSE7084, and GSE98278.

| Gene | Entrez ID | Log2FC | Ave Expr | P Value | Adj P Value | Change |
| --- | --- | --- | --- | --- | --- | --- |
| FOSB | 2354 | 3.087 | 11.371 | 1.034E-12 | 4.747E-10 | up |
| NR4A2 | 4929 | 1.810 | 9.525 | 1.410E-06 | 5.325E-05 | up |
| EGR2 | 1959 | 1.740 | 8.382 | 2.312E-06 | 7.890E-05 | up |
| CCR7 | 1236 | 1.708 | 6.920 | 1.208E-03 | 1.034E-02 | up |
| EGR1 | 1958 | 1.262 | 11.512 | 2.730E-06 | 9.038E-05 | up |
| GPR183 | 1880 | 1.208 | 8.931 | 3.414E-03 | 2.222E-02 | up |
| BANK1 | 55024 | 1.102 | 5.766 | 7.335E-04 | 7.154E-03 | up |
| FKBP11 | 51303 | 1.077 | 8.925 | 1.498E-03 | 1.213E-02 | up |
| APLNR | 187 | 1.058 | 7.632 | 1.667E-04 | 2.258E-03 | up |
| CCDC71 | 64925 | -1.115 | 4.909 | 5.570E-23 | 9.204E-19 | down |
| S1PR3 | 1903 | -1.279 | 7.328 | 1.762E-04 | 2.359E-03 | down |
| GLDN | 342035 | -1.446 | 6.657 | 8.482E-04 | 7.999E-03 | down |
| GALNT15 | 117248 | -1.642 | 6.201 | 5.382E-11 | 1.202E-08 | down |
| NPTX2 | 4885 | -1.951 | 7.003 | 1.723E-06 | 6.254E-05 | down |

Note, Ave Expr, Average expression; Adj P Value, Adjustment P Value

The expression patterns of m6A regulators were investigated based on the datasets GSE57691, GSE7084, and GSE98278, as depicted in **Figure 4E**. Our analysis revealed seven differentially expressed m6A regulators (IGF2BP1, HNRNPA2B1, YTHDC1, METTL14, YTHDC2, HNRNPC, and ZC3H13) between AAA and control tissues, which we classified as differentiated m6A regulators (DMRs). Specifically, HNRNPA2B1, YTHDC2, and HNRNPC exhibited higher expression levels in AAA tissues, whereas other regulators were down-regulated in AAA tissues compared to the control aortic tissues.

To gain further insights into the regulatory interactions between significant m6A regulators and hub DMGs, we conducted correlation analyses and presented the results in **Figure 4F**. The findings shed light on the potential regulatory associations between these m6A regulators and the identified hub genes, which play pivotal roles in AAA development and progression.

### Integrative AAA m6A-related mRNA signature (AMRMS) predicts large AAA Cases with high accuracy and highlights FKBP11 with the highest coefficient

We develop a key gene set of 14 hub DMGs which, along with 7 DMRs, compose the AAA m6A-Related mRNA Signature (AMRMS). Subsequently, we employed a comprehensive machine learning-based algorithm to conjecture whether AMRMS is a predictive risk factor for large AAA. In our analysis, two independent datasets, GSE57691 and GSE98278, were utilized, containing samples from 29 large AAAs (lAAAs) and 20 small AAAs (sAAAs), and 16 large AAAs and 15 small AAAs, respectively.

To further assess the predictive capability of AMRMS for the progression of sAAA to lAAA, we trained and tested 101 distinct predictive models through a LOOCV framework. The predictive performance of each model was then quantified using the C-index across the two datasets (GSE57691 and GSE98278). Notably, the GBM model incorporating AMRMS surpassed other models in predicting the accuracy of large AAA cases, with the highest C-index value of 0.863 (**Figure 5A**). ROC curve analysis demonstrated that AMRMS effectively differentiated between large and small AAAs, with AUC values of 1.00 in GSE57691 and 0.725 in GSE98278, respectively (**Figure 5B** and **5C)**. These results suggest that AMRMS is a robust and accurate predictor of the maximum diameter of AAAs, or in other words, a related risk factor gene set for AAA progression.

**Figure 5.**
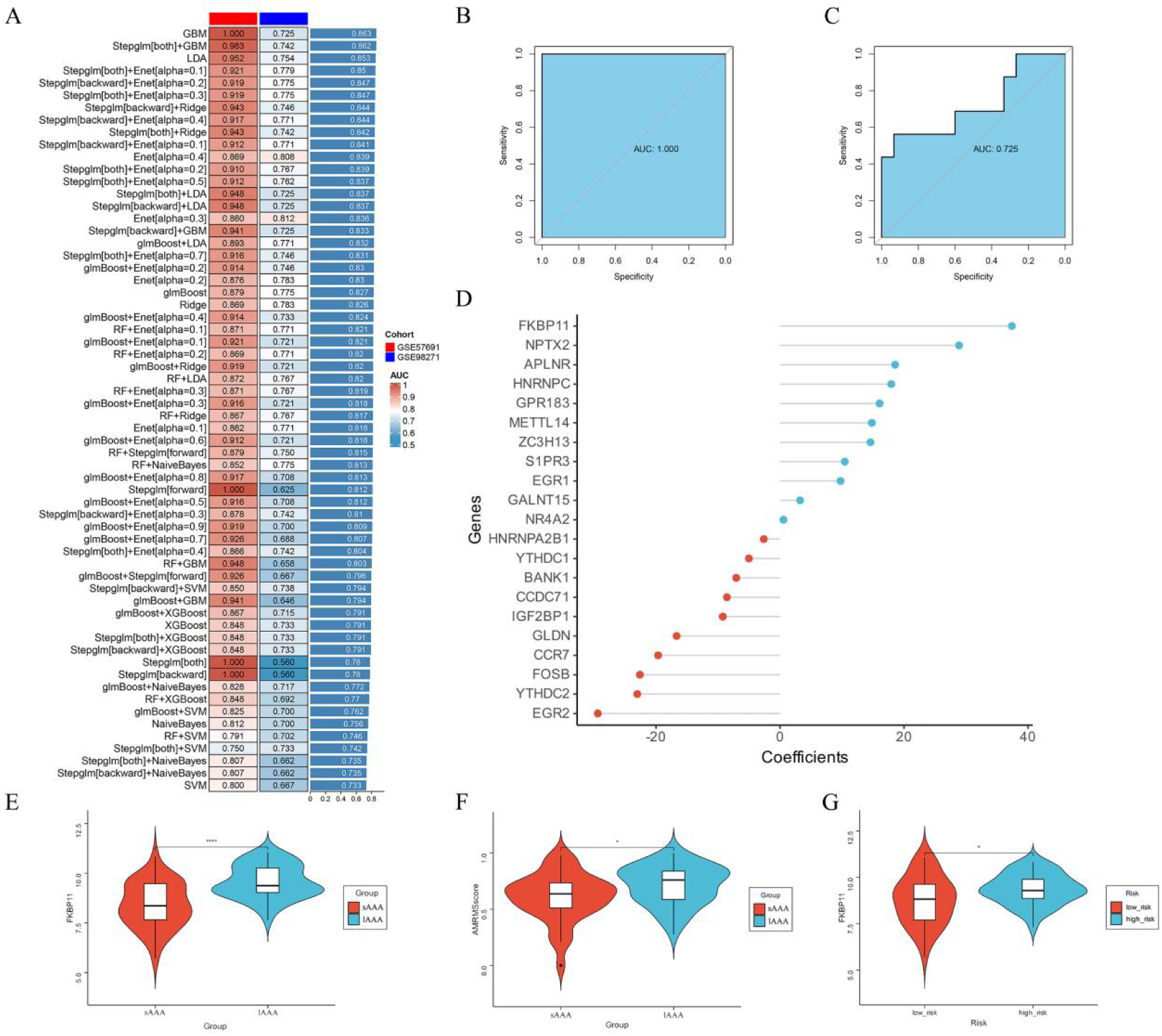
The consensus AMRMS was developed and validated to predict the risk of large AAA via the machine learning-based integrative procedure. (**A**) A total of 62 kinds of prediction models via LOOCV framework and further calculated the C-index of each model across all validation datasets. ROC curves of AMRMS to predict the risk of large AAA in GSE57691 (**B**) and GSE98278 (**C**). (**D**) Coefficients of 21 mRNAs of AMRMS finally obtained in GBM model. (**E**) The comparison of the AMRMSscore prediction model between the large AAA group and the small AAA group as shown in the violinplot. (**F**) The expressions of FKBP11 were compared between the high-risk group and the low-risk group. (**G**) The violinplot showed the expression of FKBP11 in the large AAA group compared to the small AAA group.

Furthermore, by weighting the expression levels of the 21 mRNAs enrolled in AMRMS according to their regression coefficients in the GBM model, we identified that FKBP11 was most significantly correlated with the predictive ability of AMRMS for large AAA (**Figure 5D**). We established an AMRMSscore prediction model based on the median ssGSEA score of the 21 AMRMS mRNAs, stratifying 80 patients into high and low-risk groups at the median score (50%). We observed that the AMRMSscore in large aneurysms was significantly higher than in small aneurysms, with FKBP11 expression notably higher in samples with elevated AMRMS scores (**Figure 5E** and **5F**). Furthermore, FKBP11 expression was significantly greater in the large AAA group compared to the small AAA group (**Figure 5G**). These findings underscore the pivotal role of FKBP11 in AAA progression.

### Elevated FKBP11 expression in B cell population residing in AAA samples based on scRNA-seq

ScRNA-seq was further conducted to delineate the location and functional implications of FKBP11 in AAA samples. We present baseline clinical data for six participants from our center, including four individuals with AAAs and two healthy controls (HCs), in **Table 1**. Following quality control measures, a total of 68,326 cells were profiled, consisting of 37,743 from AAA specimens and 30,583 from the healthy control specimens.

Using UMAP analysis for unsupervised clustering, we identified 18 distinct cell clusters (**Figure 6A**). These clusters were characterized based on gene expression profiles of known canonical biomarkers, from which seven cell populations were manually annotated: mononuclear phagocytes (MPs), endothelial cells (ECs), fibroblasts + mural cells (Fib_MCs), T cells + natural killer cells (TC_NKs), B cells (BCs), basophils (Baso), and plasmic dendritic cells (pDCs), visualized in a UMAP plot (**Figure 6B-6E**).

**Figure 6.**
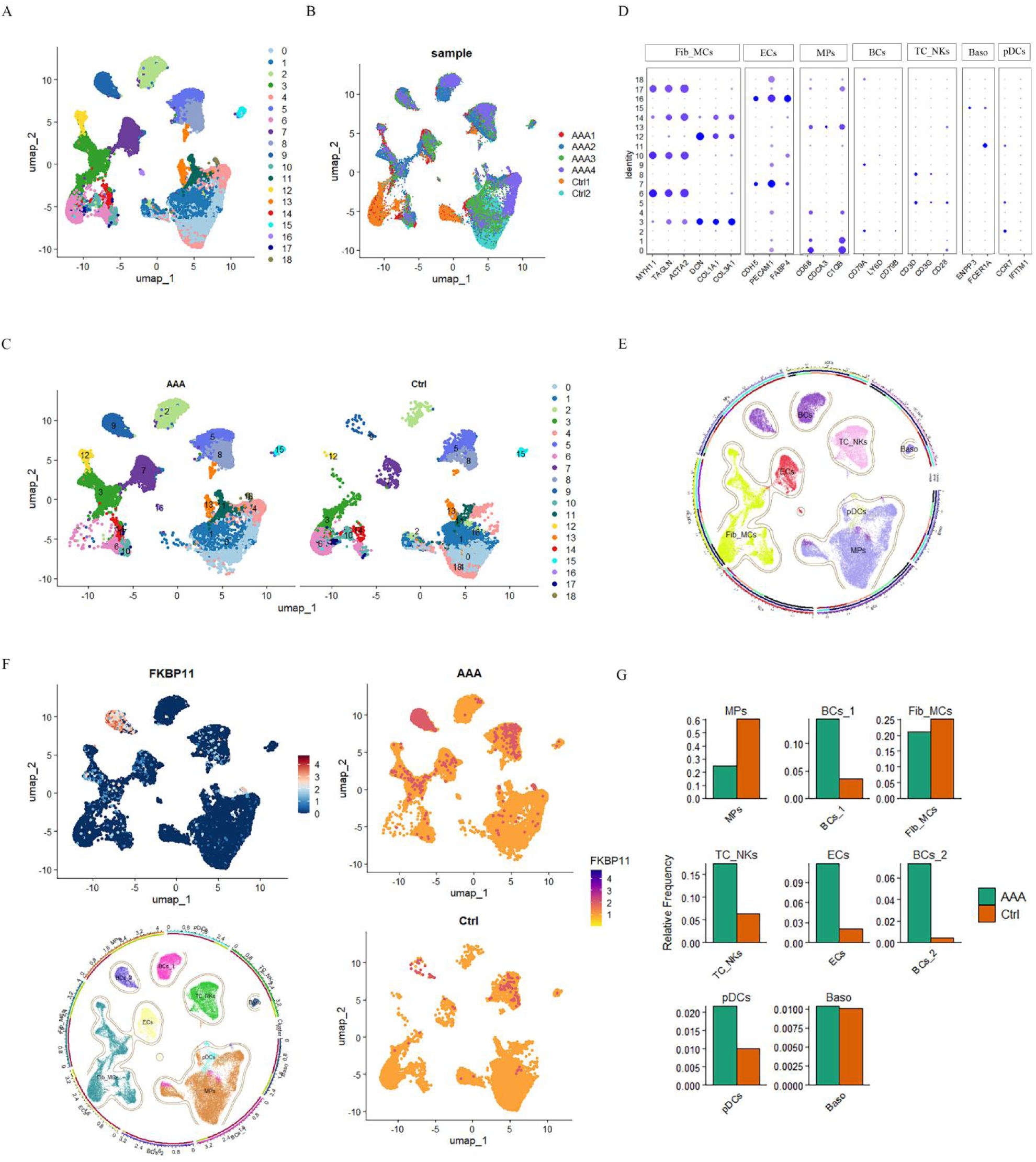
Identification of cell clusters present in the human AAA tissues and healthy aortic samples by ScRNA-seq. (**A**) The UMAP plot of all cell clusters at a resolution of 0.5 presented in all samples. (**B**) The UMAP plot of all cells according to the enrolled samples. (**C**) The UMAP plot of all cell clusters presented in the AAA groups and Ctrl group colored according to identified clusters. (**D**) The dotplot of the expressions of classical cell markers. (**E**) The UMAP plot of cell clusters presented in all aortic samples colored according to identified clusters, respectively. (**F**) The expression and location of FKBP11 was shown in the UMAP plot. (**G**) Percentages of cell populations in the AAA group and the Ctrl group.

UMAP exhibited two subclusters of BCs, and the featureplot highlighted the BCs as the predominant population of FKBP11 expression, especially within Cluster 9 and Cluster 18 (**Figure 6F**). Consequently, BCs were categorized into BCs-1 (Cluster 2), with the low level of FKBP11 expression, and BCs-2 (Clusters 9 and 18), with pronounced FKBP11 expression, designated as FKBP11*^high^* BCs.

The cell population proportions in AAA and healthy aortic tissues were noted (**Figure 6G**). Compared with the controls, the percentages of both BCs-1 and BCs-2 clusters in AAA samples were elevated. In contrast, Fib_MCs and MPs represented a smaller fraction in AAAs (21.0% and 24.9%, respectively) compared to healthy controls (25.1% and 60.4%, respectively). Furthermore, the percentages of BCs, TC_NKs, ECs, and pDCs were found to be increase in AAA samples compared to the healthy control group.

### High expression of FKBP11 in B Cells Correlates with endoplasmic reticulum stress activation

Subsequently, through AUCell score analysis, we examined 50 hallmark gene sets across various cell types (**Figure 7A**). Notably, the hallmark for the unfolded protein response was significantly upregulated in FKBP11*^high^* BCs (BC-2 Cluster) within the AAA, as compared to the control group, while it was markedly downregulated in FKBP11*^low^* B cells (BC-1 Cluster). These findings suggest that the high expression of FKBP11 is likely associated with the unfolded protein response and consequent endoplasmic reticulum (ER) stress within B cells. Further assessment with Aucell scoring of gene sets revealed the most active local activity of unfolded protein response genes within the BC-2 subgroup (**Figure 7B** and **7C**).

**Figure 7.**
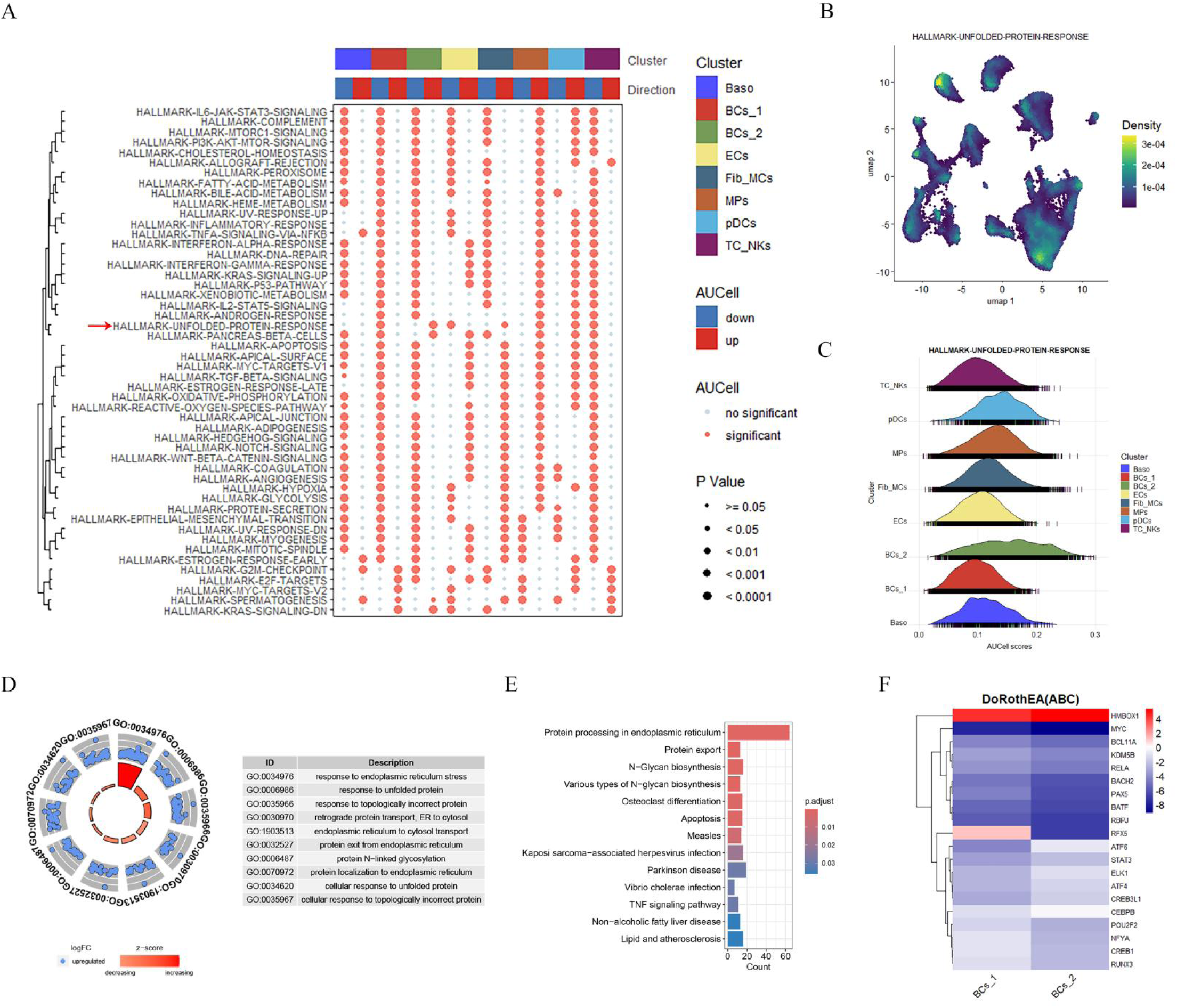
Modulation of the unfolded protein response by FKBP11 expression in B Cell subpopulation. (**A**) AUCell score analysis of 50 hallmark gene sets across different cell types between the AAA group and the Ctrl group.(**B**) Density plot of the Hallmarker unfolded protein response gene activity across cell types. (**C**) Distribution of AUCell scores for the unfolded protein response across various cell clusters. (**D**) Circos plot showing the GO enrichments of high variability genes in FKBP11^high^ BCs. (**E**) Bar graph depicting the KEGG enrichments of high variability genes in FKBP11^high^ BCs. (**F**) Heatmap of transcription factor analysis in the FKBP11^high^ BCs and FKBP11^low^ BCs.

Our analysis then delved into the high variability gene characteristics specific to FKBP11*^high^* BCs. Thus, the high variability genes of FKBP11*^high^* BCs showed significant correlations with ER functions, including “response to ER stress”, “reaction to unfolded proteins”, and “protein processing in the ER” (**Figure 7D** and **7E**). Transcription factor analysis within the BC-2 cluster identified ER function-related transcription factors such as ATF6 and ATF4 (**Figure 7F**) [11, 12].

Consequently, our analyses have uncovered a close correlation between the high expression of FKBP11 and ER stress within B cells. This association emerged as a prominent finding after machine learning analysis, which identified FKBP11 as the gene with the highest correlation to AMRMS in scRNA-seq of AAA tissues, laying the groundwork for pinpointing its expression and functional localization.

### Subgroup analysis confirms high FKBP11 expression in PCs of AAA Tissue and its correlation with ER stress

Given our discovery of high FKBP11 expression in B cells, we conducted a further subgroup analysis on our scRNA-seq data to detect in which B cell subpopulations FKBP11 was overexpressed. Unsupervised clustering divided B cells into seven subgroups (**Figure 8A-8C**). Based on the biomarkers of classic B cell subpopulations, we found that clusters 0, 1, and 3 exhibited similar marker expression, characteristic of memory B cells (Bmem), preliminarily designated as Bmem. Clusters 2, 4, 5, and 6 showed plasma cell (PC) traits and were preliminarily annotated as PCs.(**Figure 8D** and **8E**) Compared to normal aortic tissue, AAA contained a higher proportion of PC and a lower proportion of Bmem (**Figure 8F**). GSEA analysis of hallmark gene sets indicated significant upregulation of inflammation-related pathways in Bmem, such as TNF signaling, inflammatory response, IFN-γ response, and INF-α response. In PCs, pathways related to unfolded protein response and protein secretion were significantly upregulated, which may be associated with increased activation and antibody-secreting capabilities of PCs (**Figure 8G-8I**). GO and KEGG analyses of HVGs in Bmem revealed an association with ribosome activation, enriched in GO terms like cytosolic ribosome, cytoplasmic translation, and structural constituent of ribosome, and the ribosome pathway in KEGG (**Figure 8J** and **8K**). This suggests that Bmem is more involved in RNA translation processes. Conversely, the HVGs of PC cells were closely related to endoplasmic reticulum activity, enriching GO terms such as endoplasmic reticulum protein-containing complex, oxidoreduction-driven active transmembrane transporter activity, response to endoplasmic reticulum stress, and the KEGG pathway of protein processing in the endoplasmic reticulum (**Figure 8L** and **8M**). This indicates a more vigorous protein processing, transport, and metabolism function in PCs.

**Figure 8.**
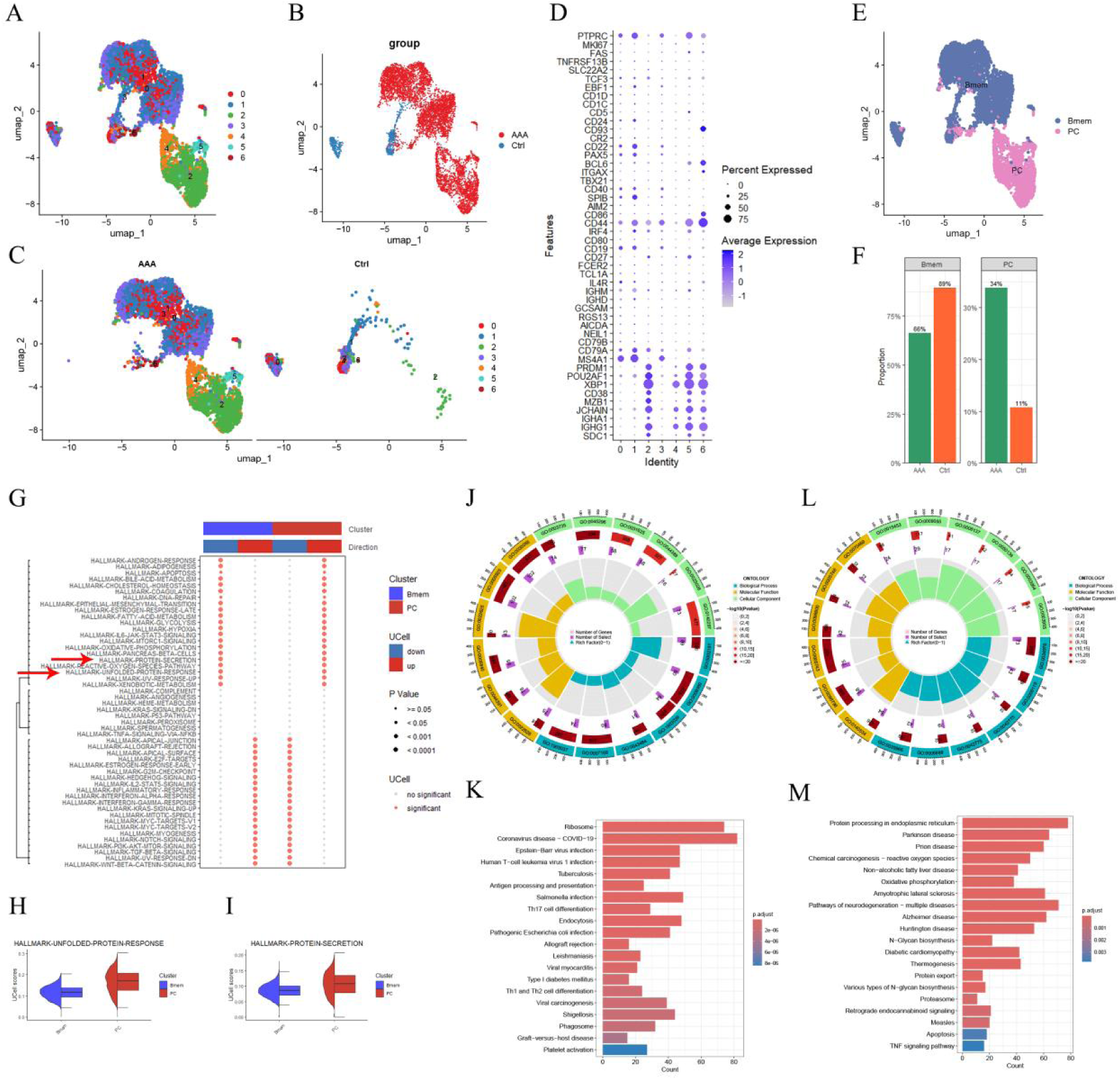
The subpopulation analysis of B cells. (**A-C**) UMAP visualization showing the unsupervised clustering of B cells into seven distinct subgroups. (**D**) Dotplot illustrating the expression of selected marker genes across the identified B cell clusters. (**E**) UMAP plot highlighting the clusters preliminarily annotated as Bmem and PCs, based on their biomarker expression profiles. (**F**) Bargraph depicting the proportion of Bmem and PCs in normal aortic tissue versus AAA. (**G**) Heatmap of the AUCell score analysis showing significant pathway enrichment in the Bmem cluster and the PC cluster. (**H-I**) Box plots representing the enrichment scores of hallmark gene sets associated with unfolded protein response and protein secretion in PCs. (**J**) GO analysis of high variability genes in Bmem. (**K**) KEGG analysis of high variability genes in Bmem. (**L**) GO analysis of high variability genes in PC. (**M**) KEGG analysis of high variability genes in PC.

Subsequent featureplot visualizations specifically demonstrated FKBP11 expression predominantly in PCs and, to a lesser extent, in Bmem (**Figure 9A** and **9B**). Violin plots revealed that FKBP11 expression in the B cell clusters of the AAA group was significantly higher than that in normal aortic B cells of the control group (**Figure 9C**). Comparisons of FKBP11 expression across different subgroups within AAA and control groups revealed significantly higher expression in both PC and B_mem subgroups within the AAA tissue (**Figure 9D**). Immunohistochemistry on consecutive sections of AAA tissue with J CHAIN as a PC marker confirmed high levels of FKBP11 localized to plasma cells within AAA tissue (**Figure 9E**). Further ultrastructural analysis via electron microscopy revealed endoplasmic reticulum expansion, autophagy, and an increase in mitochondria - characteristics of endoplasmic reticulum stress in the resident plasma cells of AAA tissue (**Figure 9F**).

**Figure 9.**
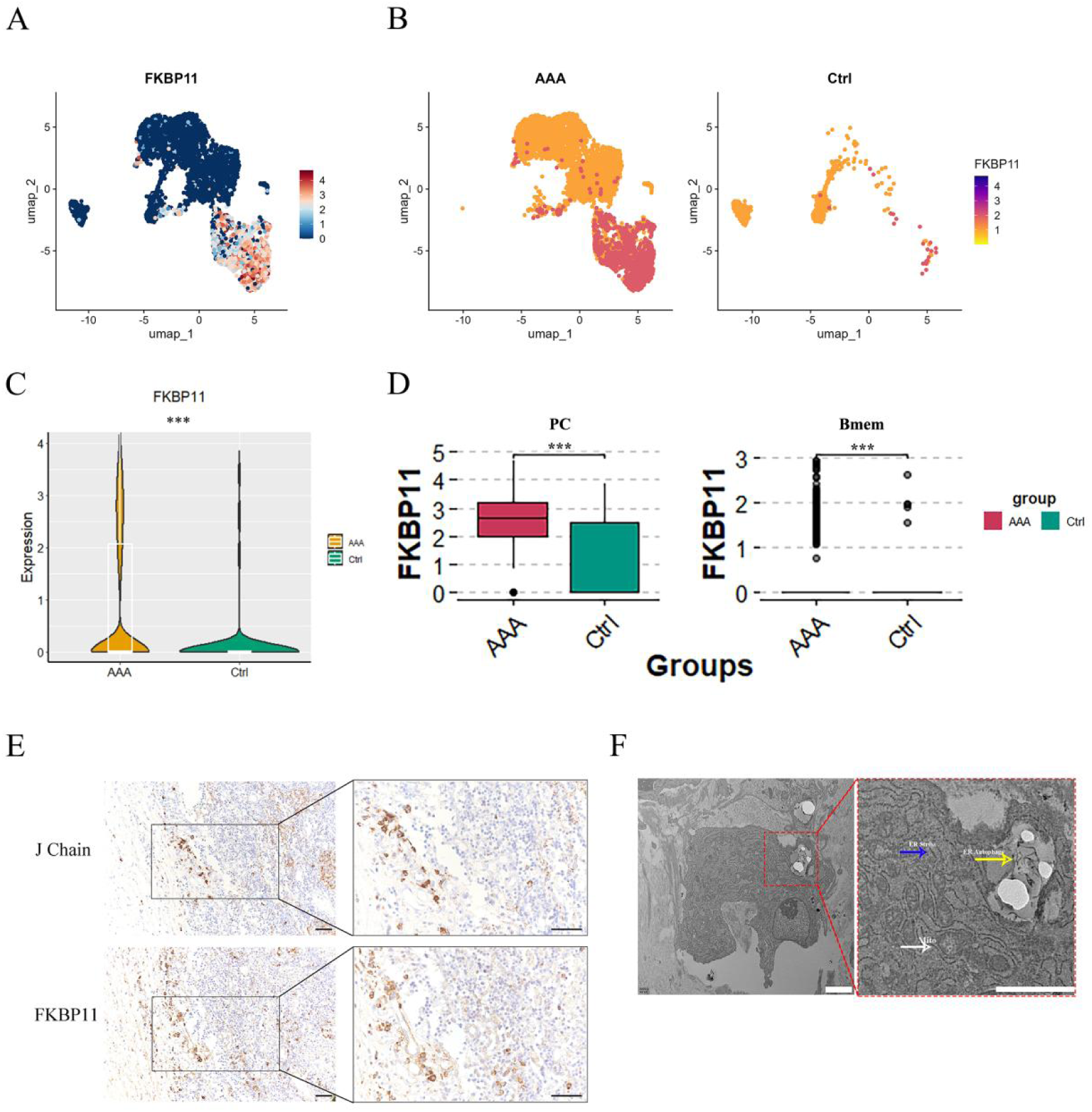
The expression and function of FKBP11 in PC. (A) UMAP plot showing FKBP11 expression levels across different subpopulation. (**B**) The respective expression of FKBP11 in the AAA group and the control group shown in UMAP plots. (**C**) Violin plot illustrating FKBP11 expression in B cell clusters. (**D**) Box plots displaying FKBP11 expression levels in PCs and Bmem within AAA and control tissues. (**E**) Reprecentative images of serial sections of human AAA tissue stained for J CHAIN and FKBP11. Scale bar, 200μm. (**F**) Electron microscopy image of AAA tissue highlighting features of endoplasmic reticulum stress in plasma cells, including endoplasmic reticulum expansion (blue arrow), autophagy (yellow arrow), and increased mitochondria (white arrows).

Together with the above results and the biological function of FKBP11 as a key enzyme in ER stress [13], we postulate that the high expression of FKBP11 in resident plasma cells within AAA tissue correlates with ER stress.

### Pseudotime analysis reveals FKBP11’s involvement in the differentiation from Bmem to PC

Cytotrace analysis was employed to assess the stemness and differentiation potential of B cell subgroups. As depicted in **Figure 10A-10C**, Bmem cluster of the AAA group displayed higher Cytotrace scores, predominantly in the range of 0.5-0.9, indicating substantial differentiation potential. Conversely, PC cluster in the AAA group primarily exhibited lower Cytotrace scores, concentrated between 0-0.4, suggesting they are at a later stage of differentiation compared to AAA group Bmem. For the control group, both Bmem and PC within normal aortic tissue samples presented with lower differentiation potential, potentially indicative of the unactivated status of B cell subpopulations. Correlation analysis between gene expression and Cytotrace scores revealed a negative association with FKBP11, positing that cells with elevated FKBP11 expression are likely in the terminal phase of differentiation (**Figure 10D** and **10E**).

**Figure 10.**
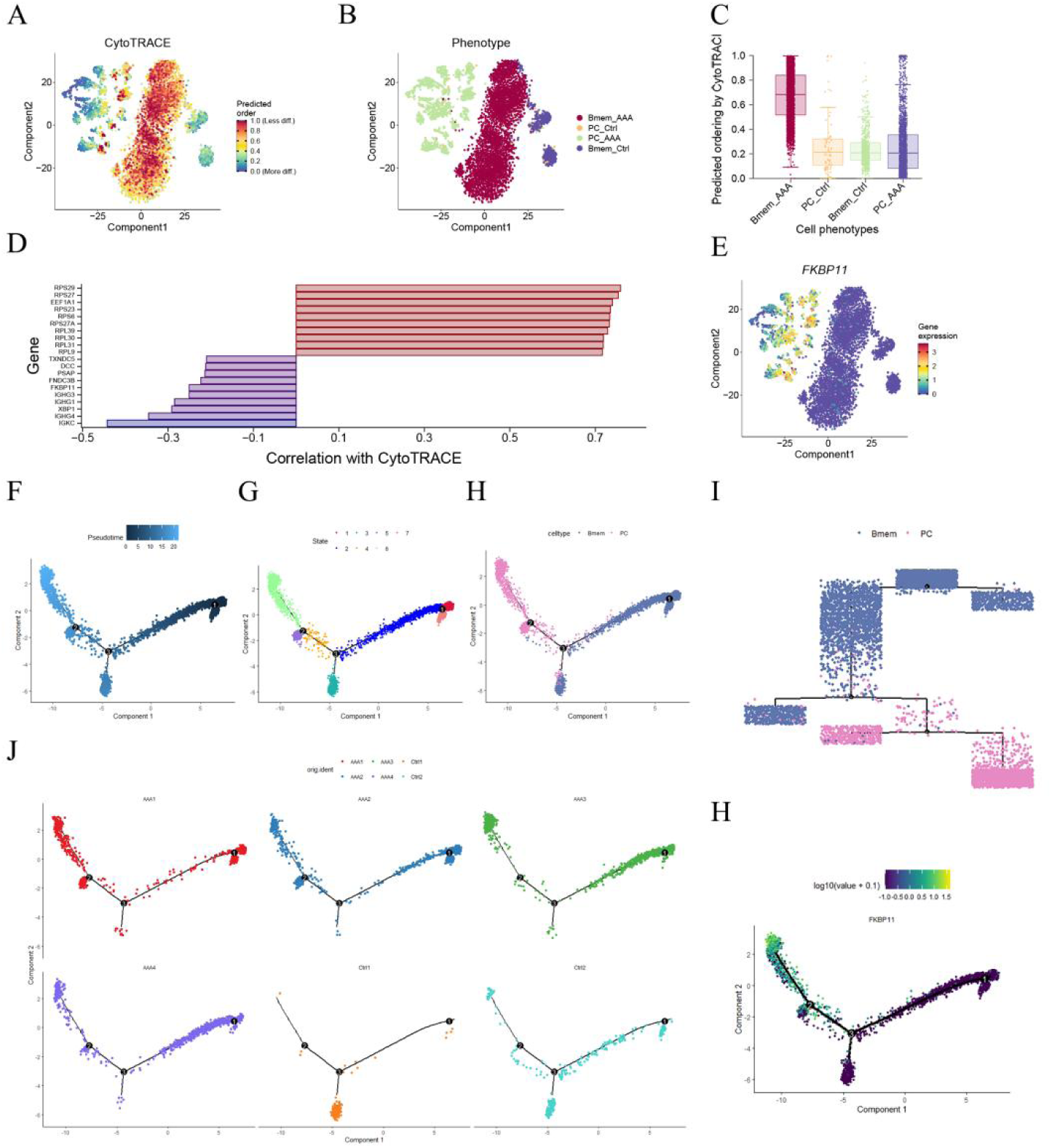
The Cytotrace analysis and pseudotime analysis of B cell subpopulations. (**A, B**) t-Distributed Stochastic Neighbor Embedding (tSNE) plot of B cell subpopulations from AAA and Ctrl depicting the distribution of CytoTRACE scores among B cells. (**C**) Box plot representing the distribution of predicted stemness by Cytotrace across different B cell subphenotypes from AAA and Ctrl. (**D**) Bar graph depicting the correlation of Top 20 genes with Cytotrace scores. (**E**)The t-SNE plot show the correlation of FKBP11 gene expression and B cell subpopulations from AAA and Ctrl. (**F**) The single-cell trajectory of B cell subpopulations predicted by Monocle 2 and ordered in pseudotime. (**G**) The distribution of B cells in the pseudotime trajector colored according to 7 distinct states. (**H**, **I**) The distribution of B cells in the pseudotime trajector colored according to identified subclusters. (**J**) Trajectory plots for individual samples from the AAA and Ctrl groups, displaying the progression of B cells along the main trajectory and branches. (**K**) The distribution of FKBP11 gene expression along the main trajectory and branches.

To further elucidate the role of FKBP11 in the differentiation process among B cell subpopulations, pseudotime analysis was conducted using the Monocle 2R package. B cells were mapped on a pseudotime trajectory created by the analysis, divided into seven distinct states, comprising a main trajectory (states 1, 2, 4, 6) and three branches (states 3, 5, 7) (**Figure 10F** and **10G**). Bmem were principally situated at the onset of the main trajectory, while PCs were localized predominantly towards the latter half of the trajectory, illustrating the transformation from Bmem to PC within the BC cell clusters (**Figure 10H** and **10I**). In the AAA group, B cells from four samples were mainly positioned along the main trajectory, transitioning from Bmem to PC cells. In contrast, B cells in the control group were primarily distributed in the main trajectory’s states 1 and 2 and the branch state 3, with a discernible gap between the main and branching trajectories (**Figure 10J**). Subsequent findings showed an upregulation of FKBP11 expression concurrent with the shift from Bmem to PC, underscoring FKBP11’s significant role in the activation and differentiation of B cells into plasma cells (**Figure 10H**).

Our results provide a nuanced understanding of the molecular mechanisms underlying B cell differentiation, highlighting FKBP11 as a key regulator.

### Correlation between elevated FKBP11 expression and m6A expression patterns in PCs within AAA walls

Through scRNA-seq analysis, we have discerned a close association between high FKBP11 expression and ER stress in PCs, as well as B cell differentiation. We further analyzed the correlation between FKBP11 and m6A methylation within PCs of AAA samples utilizing scRNA data (**Figure 11A**-**11C**). Initially, m6A scores were assigned to different cells in both the AAA and control groups, based on the expression levels of 21 m6A modulators mentioned before. We observed that within all clusters, the BC-2 cluster with high FKBP11 expression exhibited the highest m6A scores, and significantly surpassing the m6A scores in the BC-1 cluster with lower FKBP11 expression. Notably, in the AAA group, the BC-2 cluster with high FKBP11 expression had a significantly higher m6A score compared to the BC-1 cluster, a trend similarly reflected in the Ctrl group.

**Figure 11.**
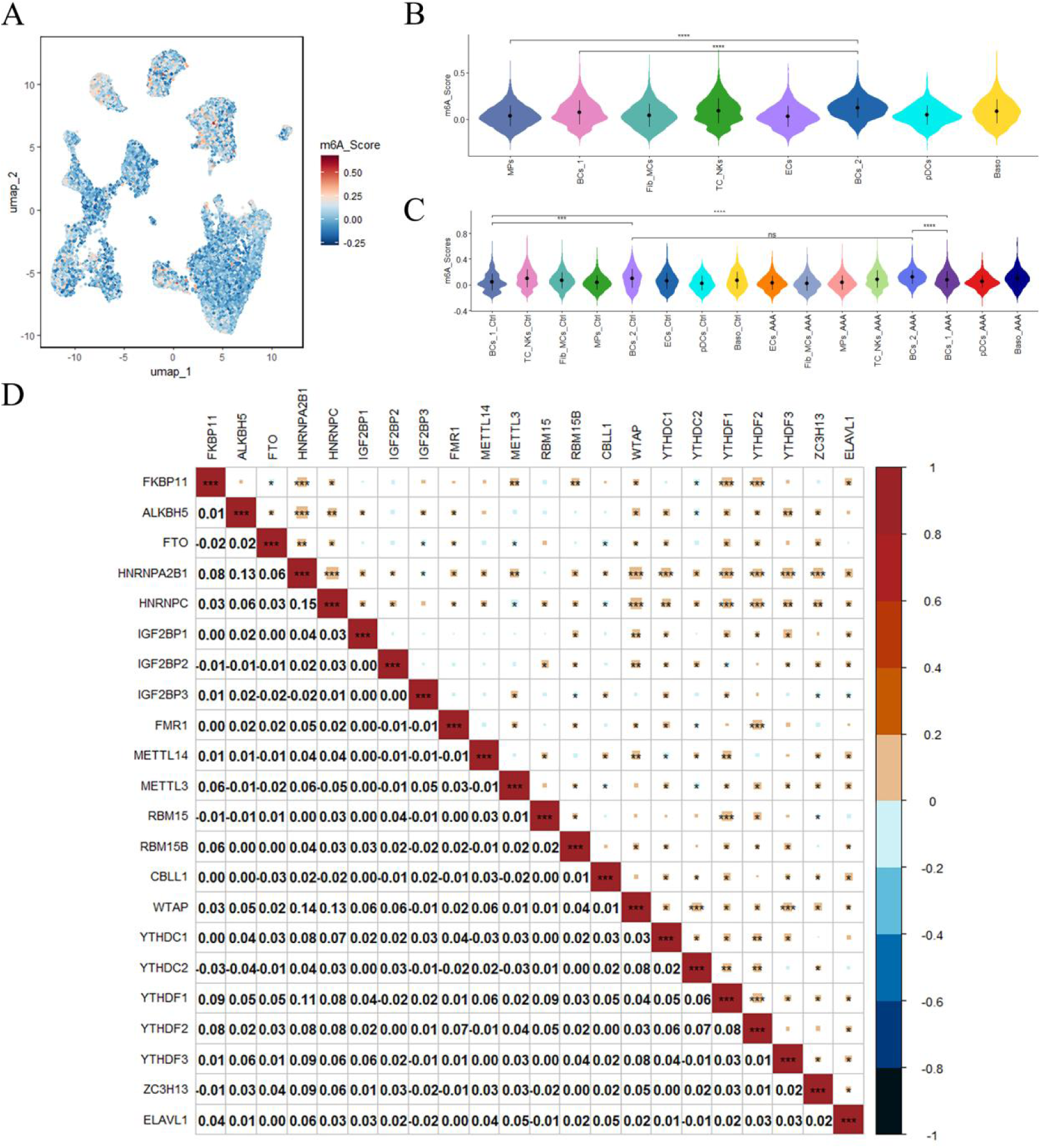
Association of FKBP11 expression with m6A methylation patterns in PCs from AAA walls. (**A**) UMAP plot visualizing m6A scores across different cell populations in AAA and control groups, with color intensity indicating the degree of m6A methylation. (**B**) Violin plots depicting the m6A scores across different clusters. (**C**) Violin plots contrasting m6A scores in the AAA and control groups. (**D**) Heatmap of correlation coefficients between FKBP11 expression and 21 m6A modulators in the BC-2 cluster of PCs from AAA walls. The color scale reflects the strength of positive (red) to negative (blue) correlations, with asterisks indicating levels of significance, *p < 0.05, **p < 0.01, ***p < 0.001.

Subsequent correlation analysis between FKBP11 expression and m6A modulators within the BC-2 cluster of the AAA group was performed (**Figure 11D**). Specifically, FKBP11 gene expression showed a significant positive correlation with key m6A writers and readers, including METTL3, WTAP, RBM15B, YTHDF1, YTHDF2, HNRNPC, and HNRNPA2B1. Additionally, FKBP11 expression was found to be negatively related to the m6A demethylase FTO and the m6A reader YTHDC2.

## Discussion

AAA with its high incidence and mortality rates, poses a significant clinical challenge, especially in managing patients with small and asymptomatic AAAs that do not meet the criteria for surgical intervention. The unpredictability of enlargement and rupture of these AAAs necessitates a fundamental investigation and definition of their cellular and molecular nature. Our study highlights the critical role of m6A methylation in the pathogenesis of AAA, identifies it as a promising therapeutic target. Utilizing MeRIP-seq, we have uncovered a distinct RNA m6A modification landscape in AAA tissues compared to healthy controls, offering new insights into how m6A is involved in AAA development. Furthermore, combining RNA-seq data with bioinformatics and machine learning, we identified m6A-related genes, AMRMS, that could potentially predict the progression of AAA from small to large and highlight the role of FKBP11 in the prediction. Notably, through scRNA-seq, we have detected FKBP11, revealing the potential mechanism by which m6A-FKBP11 exacerbates AAA through the regulation of ER stress in PCs. This research, integrating multi-omics analysis and machine learning, for the first time, underscores the complex relationship between m6A modification and AAA progression, opening new avenues for targeted therapeutic strategies based on molecular insights.

We initially performed MeRIP-seq as the principal method to conduct a comprehensive analysis of m6A modifications in AAA tissue samples, comparing to healthy control aortic vessels. We identified overlapping functions and signaling pathways involved in m6A modifications between the two sample groups, such as “ubiquitin-mediated proteolysis”, “TGF-β signaling pathway”, and “protein processing in the endoplasmic reticulum”, illustrating the conserved nature of m6A regulation within the aortic wall across different states. However, the differences in specific modified mRNAs and the extent of methylation modifications between the two groups significantly influence cellular function differentiation within aortic tissues. This diversity outlines m6A methylation as a complex, dynamic regulatory network. Remarkably, mRNAs with differential m6A modifications were significantly enriched in pathways directly associated with AAA development and progression, such as the TGF-β signaling pathway, smooth muscle cell contraction, arteriosclerosis, and autophagy. These findings emphasize the complex role of m6A in controlling the post-transcriptional fate of key mRNAs, thereby affecting the disease trajectory of human AAA. This nuanced interplay between the conserved and differential aspects of m6A modifications enhances our understanding of AAA’s molecular mechanisms, demonstrating how subtle shifts in mRNA destiny can lead to significant changes in AAA disease outcomes.

Subsequently, we embarked on the development of a novel m6A-related gene signature AMRMS to forecast the progression of AAA. Previous studies have employed similar bioinformatics approaches to investigate the effects of m6A methylation on the progression or prognosis of AAA [9, 14–17]. However, these studies only analyzed the expression patterns of m6A modulators, but overlooked the analysis of m6A-modified RNAs, rendering their models for predicting the impact of m6A on AAA incompletely.

Our initiative diverged by not only assessing the differential expression of m6A-modified RNAs but also by scrutinizing the expression variances of m6A modulators. Leveraging our analytical insights, we curated and constructed the geneset AMRMS, which, through machine learning techniques, demonstrated robust efficacy in differentiating between sAAAs and lAAAs, serving as a risk marker for the transition from sAAAs to lAAAs. The aggregate metrics, including the ROC curve’s AUC and the C-index, surpassed a previous model [18], offering enhanced specificity in forecasting AAA progression. The AMRMS, the gene signature we developed, stands as an advanced m6A-associated AAA risk prediction model. This advancement enriches the bioinformatics and machine learning framework for understanding AAA, contributing a novel tool for risk stratification and tailored management strategies. Consequently, it may lay the groundwork for devising efficacious targeted interventions aimed at halting AAA progression and ameliorating patient prognoses.

In AMRMS, the FKBP11 gene exhibited the highest regression coefficient, playing a pivotal role in this predictive model. Consequently, further analysis was conducted on FKBP11 in our study. Given the complex cellular composition of AAA, scRNA-seq offers an unbiased view into the molecular spectra and the heterogeneity of cellular components within the AAA vascular wall, providing crucial insights into AAA research. Our study, leveraging single-center scRNA data from our institution’s vascular surgery department, delved into the expression localization and function of FKBP11. We discovered that FKBP11 was specifically expressed in the PC subpopulation within the BC population. Despite identifying a significant presence of PCs in AAA walls through pathological sections, previous studies reporting an increase in the BC/PC population in human AAA, and the removal of B cells has been shown to have a protective effect in AAA animal models [19, 20]. However, primary PCs within the aneurysmal wall are challenging to isolate and have a short lifespan ex vivo, meanwhile the PCs induced from peripheral blood B cells and commercial plasma cell lines hardly mimic the complex vascular environment in the AAA walls, making the role of AAA-resident PCs in AAA progression elusive. Integrating machine learning outcomes with scRNA-seq data, we hypothesize that high expression of FKBP11 in PCs is a risk factor for AAA progression. The up-regulation of FKBP11 might promote the progression of AAA by activating ER stress in plasma cells. FKBP11 is a critical enzyme regulating the UPR on the endoplasmic reticulum, potentially reshaping the UPR signaling pathway to participate in various diseases. In PCs, FKBP11 acts as a specific antibody folding catalyst upregulated during ER stress [21, 22]. It modulates the UPR to accommodate the increased load of nascent immunoglobulins and prevent the accumulation of misfolded proteins in the ER [23, 24]. Increasing evidence suggests that human AAA exhibits significant autoimmune characteristics, including the secretion of autoantibodies and the dysfunction of regulatory T cells, which are believed to promote AAA progression [25–28]. In our study, pseudotime analysis indicates that PCs in the AAA cell wall originate from memory B cells, during which FKBP11 expression is elevated. Through electron microscopy, we also observed PCs in the AAA walls, characterized by ER expansion, an increase in mitochondria, and signs of ER autophagy, indicating an active state of antibody synthesis and secretion under ER stress in PCss within the AAA wall. Combining the above results, a high level of FKBP11-mediated ER stress in PCs is likely a risk factor that promotes AAA progression. FKBP11 in PCs could be a potential important target for inhibiting AAA from an immunological perspective in the future.

In our final analysis, we examined the expression patterns of m6A modulators across different cell populations in the scRNA-seq data, assigning scores to each. Notably, PCs exhibited the highest m6A scores among all cell populations, with a significant increase compared to the FKBP11*^low^* BC group. This finding suggests that m6A methylation modifications are most active in PCs, and the differentiation of Bmem into PCs is likely regulated by m6A methylation modifications. Subsequently, we conducted a correlation analysis between the expression of FKBP11 and m6A modulators in PCs. Our results revealed that FKBP11 expression is significantly positively correlated with the m6A methyltransferase complex components METTL3 and WTAP, and negatively correlated with the demethylase FTO. Furthermore, FKBP11 also showed significant correlations with multiple m6A reader proteins. These findings suggest that the increased expression of FKBP11 in PCs is closely associated with m6A methylation, which likely serves as a crucial post-transcriptional regulatory target for the activation of ER stress in PCs. Currently, several m6A methylation inhibitors are in the preclinical trial phase [29]. If future developments can lead to the creation of a plasma cell-targeted drug delivery system capable of directing m6A inhibitors specifically to the activated plasma cells within the aortic walls of AAA patients, it could potentially inhibit AAA progression. Our research provides novel insights in early drug intervention and the delay of AAA progression.

Recognizing the limitations of this study is crucial. Our sample size was relatively small, including both MeRIP-seq and scRNA-seq. While our research provides evidence of m6A methylation’s involvement in AAA by focusing on the role of m6A-related mRNAs in the progression from small to large AAA, the vast amount of data means that understanding how m6A methylation affects the fate and function of RNAs in different cells, thereby contributing to AAA progression, requires extensive future efforts. Additionally, our study suggests that m6A modifications may influence ER stress in plasma cells, but the scarcity of human AAA clinical samples makes extracting and culturing a sufficient number of primary resident PCs challenging. Advances in technology in the future could aid in further exploring these mechanisms more comprehensively.

In conclusion, our research identifies the significance of the m6A methylation regulatory network in AAA pathogenesis and reveals a detailed transcriptomic profile of m6A modifications in AAA tissues. By employing bioinformatics and machine learning algorithms, we develop a novel efficient m6A-related marker, AMRMS, to predict AAA progression which highlights FKBP11 with the highest coefficient. Furthermore, our integration of scRNA-seq data suggests that m6A methylation may influence AAA progression by up-regulating the expression of FKBP11 and further affecting ER stress of plasma cells within AAA walls. These results could potentially influence AAA diagnostic and therapeutic strategies to prevent aneurysm expansion in near future.

## Abbreviations

AAA: abdominal aortic aneurysm
AMRMS: AAA m6A-related mRNA signature
AUC: Area Under the Curve
BP: biological process
CBBs: Cell Barcoded Magnetic Beads
CC: cellular component
CCA: Canonical Correlation Analysis
CMUaB: CMU Aneurysm Biobank
CDS: coding sequence
CTA: computed tomography angiography
DEGs: differential expressed genes
DMGs: differentially methylated genes
ECs: endothelial cells
Enet: Elastic Network
ER: endoplasmic reticulum
FC: Fold Change
GBM: Generalized Boosted Regression Modeling
glmBoost: Generalized Linear Model Boosting
GO: gene ontology
HC: healthy control
IHC: Immunohistochemistry
IP: immunoprecipitation
KEGG: Kyoto encyclopedia of genes and genomes
LDA: Linear Discriminant Analysis
LOOCV: Leave-One-Out Cross-Validation
m6A: N6-methyladenosine
MASS: Multicenter Aneurysm Screening Study
MeRIP-seq: methylated RNA immunoprecipitation with next-generation sequencing
MF: molecular function
MPs: mononuclear phagocytes
NKCs/TCs: natural killer cells/T cells
PCs: plasma cells
PCA: principal Component Analysis
PPI: Protein-Protein Interactions
RF: Random Forest
ROC: Receiver Operating Characteristic
SMCs: smooth muscle cells
SMGs: specific methylated genes
Stepglm: Step Generalized Linear Model
SVM: Support Vector Machine
t-SNE: t-Distributed Stochastic Neighbor Embedding
Teff: Effector T cells
Th: Helper T cells
Treg: Regulatory T cells
UMI: unique molecular identifiers
UPR: unfolded protein response
XGBoost: eXtreme Gradient Boosting
3’UTR: 3’ untranslated region
5’UTR: 5’ untranslated region.

## Authors’ contributions

YCH, and JX performed experiments and data analysis. YCH and JZ wrote the article. YL provided technical support. YSH, PE, SYW, and HJ contributed to the discussion of the project and article. JZ, BD and PE did critical editing. JZ and YCH designed research and discussed results.

